# Convergent recognition of tobamovirus movement proteins by phylogenetically distinct NLR immune receptors

**DOI:** 10.64898/2026.10.05.756766

**Authors:** Yu Sugihara, Hagit Hak, Nutthalak Laksanavilat, Juan M. Azcue, Ziv Spiegelman, Jogi Madhuprakash

## Abstract

- Tomato brown rugose fruit virus (ToBRFV) has emerged as a major threat to tomato production worldwide, overcoming the widely deployed Tm-2^2^-mediated resistance and challenging current strategies for tobamovirus disease control. The coiled-coil nucleotide-binding leucine-rich repeat (CC-NLR) immune receptor HREZ from the wild tomato *Solanum habrochaites* confers resistance to ToBRFV, but its activation mechanism remains unclear.
- We co-expressed ToBRFV proteins with HREZ in *Nicotiana benthamiana* and assessed cell death and ion leakage, examined receptor assembly by blue native polyacrylamide gel electrophoresis, predicted oligomeric states using AlphaFold 3, and reconstructed phylogenies of 4,657 plant NLR NB-ARC domains and 32 Virgaviridae movement proteins (MPs).
- HREZ is activated by the ToBRFV MP and induces hypersensitive cell death. Upon activation, it assembles into a higher-order complex, with AlphaFold 3 modelling supporting a pentameric resistosome. HREZ falls in a CC-NLR lineage distinct from the canonical tobamovirus resistance receptor Tm-2^2^. Across eight *Virgaviridae* MPs, HREZ and Tm-2^2^ have distinct recognition spectra, and the four MPs activating HREZ also promote its higher-order assembly.
- These findings reveal that phylogenetically distinct CC-NLRs can converge on tobamovirus MPs as a common recognition target while retaining distinct recognition specificities, and provide a foundation for developing resistance strategies to current and emerging tobamoviruses.

## Introduction

Plant immune receptors can independently evolve to recognize the same pathogen molecules or activities, providing striking examples of convergent evolution in host immunity. Such convergent recognition has been described in several plant immune receptor systems and demonstrates that distinct receptor lineages can independently target the same pathogen molecules (Ashfield *et al*., 2004; Carter *et al*., 2019; Ahn *et al*., 2023a; Chen *et al*., 2023; Kim *et al*., 2023). However, the evolutionary and functional consequences of such convergence remain poorly understood, including whether convergence on a shared target produces similar or distinct recognition breadth. Antiviral immune receptors provide a useful system for studying such convergence because essential viral proteins may be constrained in their ability to evade immune recognition (Hak *et al*., 2023).

Plants rely on intracellular nucleotide-binding leucine-rich repeat (NLR) immune receptors to detect pathogen invasion and activate immune responses, often culminating in localized hypersensitive cell death that restricts pathogen spread (Duxbury *et al*., 2021; Contreras *et al*., 2023a). Recent structural studies have fundamentally reshaped our understanding of NLR function by revealing that receptor activation is coupled to the assembly of higher-order oligomeric complexes, termed resistosomes, which function as signalling complexes and, in several cases, calcium-permeable channels that initiate immune responses (Wang *et al*., 2019; Zhao *et al*., 2022; Förderer *et al*., 2022; Liu *et al*., 2024; Selvaraj *et al*., 2024; Madhuprakash *et al*., 2024; Toghani *et al*., 2026b; Guo *et al*., 2026; Seager *et al*., 2026). While substantial progress has been made in defining the structural basis of NLR activation in immunity to bacterial, fungal, and oomycete pathogens, less is known about how antiviral plant NLRs respond to viral proteins and whether they employ similar resistosome-mediated mechanisms.

Plants have evolved antiviral NLRs capable of recognizing a remarkably diverse range of viral proteins, including coat proteins (CPs), movement proteins (MPs), replicases and other viral effectors. Among tobamoviruses, several distinct NLR systems recognize different viral proteins. N recognizes the Tobacco mosaic virus (TMV) replicase helicase domain (p50) (Erickson *et al*., 1999), whereas N′ and the pepper L alleles recognize tobamovirus CPs (Gilardi *et al*., 2004; Tomita *et al*., 2011; Sekine *et al*., 2012). Notably, the L alleles confer hierarchical resistance spectra against different tobamovirus pathotypes (Tomita *et al*., 2011). In contrast, Tm-2^2^ recognizes tobamovirus MPs (Weber & Pfitzner, 1998; Chen *et al*., 2017). Other well-characterized antiviral NLRs include Rx, which detects the Potato virus X CP (Bendahmane *et al*., 1995); Sw-5b, which recognizes the Tomato spotted wilt virus MP NSm (Zhu *et al*., 2017); and Ry_sto_, which recognizes the Potato virus Y CP (Grech-Baran *et al*., 2020; Grech-Baran *et al*., 2022). Despite this diversity of recognition specificities, little is known about how antiviral NLRs are activated by viral proteins and whether they employ conserved resistosome-mediated mechanisms.

Tomato brown rugose fruit virus (ToBRFV, *Tobamovirus fructirugosum*), a member of the genus *Tobamovirus* within the family *Virgaviridae*, has emerged over the past decade as one of the most serious threats to global tomato production (Salem *et al*., 2016, 2023; Zhang *et al*., 2022). Since its first detection, ToBRFV has spread rapidly across major tomato-growing regions worldwide (Salem *et al*., 2023; de Koning *et al*., 2025), causing substantial economic losses through reduced yield and fruit quality. The global impact of this virus largely reflects its ability to overcome the widely deployed *Tm-2*^*2*^ resistance gene, which had provided durable protection against tobamoviruses, including Tomato mosaic virus (ToMV), for over six decades (Lanfermeijer *et al*., 2003; Luria *et al*., 2017; Hak *et al*., 2023). Unlike other tomato-infecting tobamoviruses, ToBRFV carries a MP that evades *Tm-2*^*2*^-mediated resistance while maintaining the viral movement function required for infection (Yan *et al*., 2021; Hak & Spiegelman, 2021; Hak *et al*., 2023). Several sources of ToBRFV resistance have been identified in *Solanum* germplasm (Jewehan *et al*., 2022b,c; Zinger *et al*., 2025; Jaiswal *et al*., 2025; Zois *et al*., 2025). One such source is *HREZ* (High Resistance Enza Zaden), a novel resistance gene identified in the wild tomato species *S. habrochaites* and shown to confer high resistance to ToBRFV (Verweij *et al*., 2026). *HREZ* encodes a coiled-coil-type NLR (CC-NLR) immune receptor, as does *Tm-2*^*2*^. Previous genetic and virological studies implicated the ToBRFV MP in *HREZ*-mediated resistance, as resistance-breaking ToBRFV variants carry amino acid substitutions in the MP (Jewehan *et al*., 2022a; Zisi *et al*., 2024; Verweij *et al*., 2026). However, direct experimental evidence that HREZ responds to the ToBRFV MP has been lacking, and the evolutionary relationship between HREZ and Tm-2^2^ remains unresolved.

In this study, we investigated the activation mechanism underlying *HREZ*-mediated resistance to ToBRFV. We demonstrate that HREZ is activated by the ToBRFV MP and triggers hypersensitive cell death. Activation of HREZ promotes higher-order oligomeric assembly consistent with resistosome formation, although our biochemical analyses provide no evidence that the MP is stably retained within the assembled HREZ complex. Phylogenetic analyses further reveal that HREZ and Tm-2^2^ belong to distinct CC-NLR lineages, while comparative assays uncover distinct, partially overlapping recognition spectra for tobamovirus MPs. Together, these findings show that phylogenetically distinct CC-NLRs can converge on the same viral target without converging on the same recognition spectrum, and link HREZ recognition specificity to activation-dependent higher-order assembly.

## Results

### HREZ is activated by the ToBRFV movement protein

To determine which ToBRFV protein activates HREZ, we took advantage of the *Nicotiana benthamiana* experimental system to transiently express HREZ with ToBRFV proteins. We cloned the ToBRFV-encoded RNA-dependent RNA polymerase (RdRP), MP, and CP, and co-expressed each protein with either HREZ or Pikobody^α-GFP^, a negative-control engineered receptor that recognizes GFP (Kourelis *et al*., 2023) (**Fig. 1a,b**). Among the proteins tested, only the ToBRFV MP triggered HREZ-dependent cell death (**Fig. 1a,b**), suggesting that the ToBRFV MP acts as the avirulence determinant that activates HREZ.

**Fig. 1.**
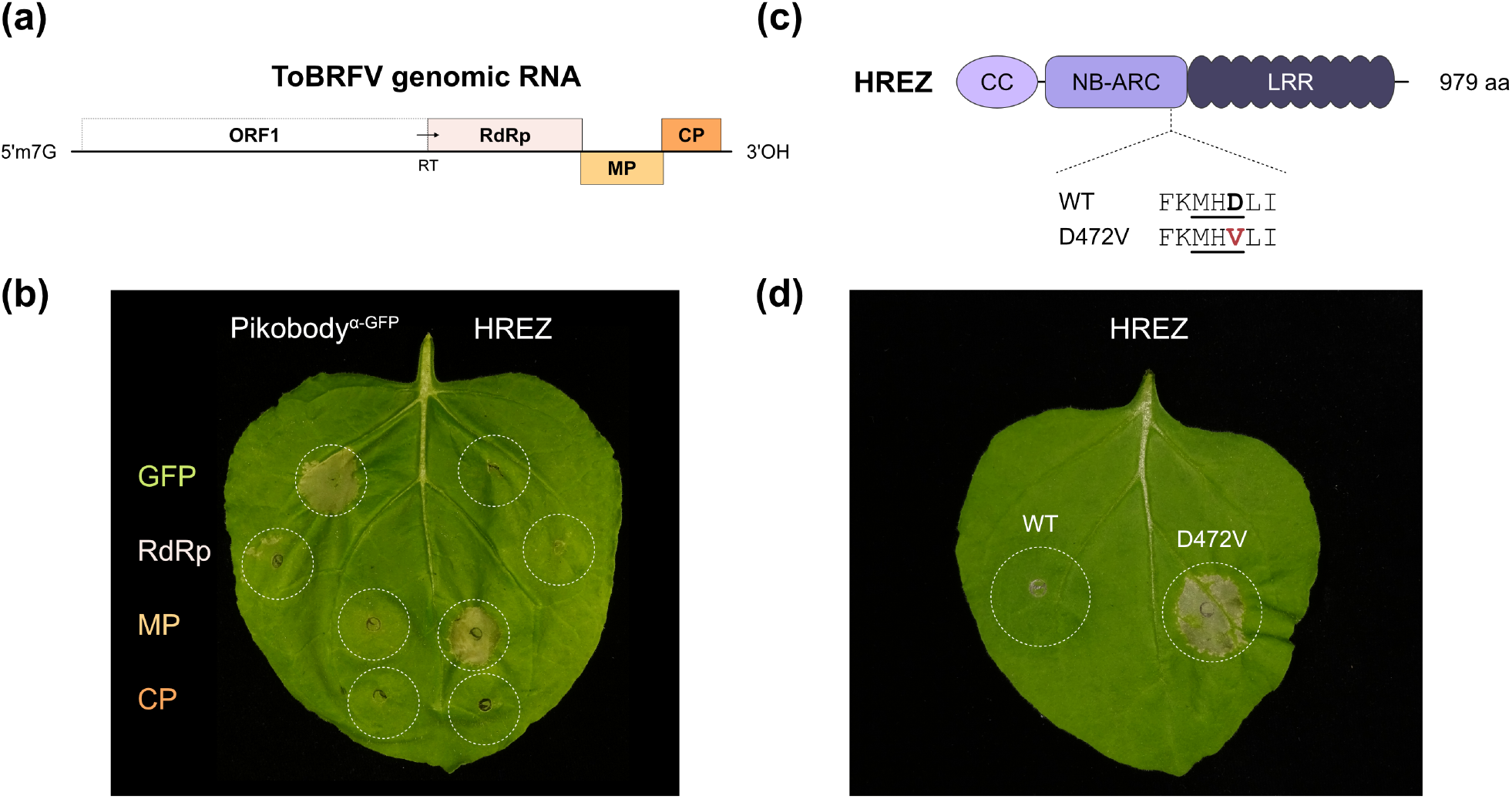
The CC-type NLR immune receptor HREZ is activated by the ToBRFV MP. (a) Schematic representation of the tomato brown rugose fruit virus (ToBRFV) genome organization and the virus-encoded proteins tested with HREZ. The genome schematic was adapted from Salem *et al*. (2023). The ToBRFV proteins tested here were based on sequences from the Tom1-Jo isolate (Salem *et al*., 2016). The ToBRFV genomic RNA contains open reading frame (ORF) 1 and regions encoding the RNA-dependent RNA polymerase (RdRp), movement protein (MP), and coat protein (CP). RdRp is produced by translational readthrough (RT) of ORF1. (b) Representative image of a cell death assay after co-expression of GFP or ToBRFV-encoded proteins with either the GFP-responsive Pikobody^α-GFP^ or HREZ. Leaves were photographed 5 days after infiltration. (c) Schematic representation of the domain architecture of HREZ and the D472V mutation in the MHD motif. HREZ consists of CC, NB-ARC, and LRR domains. (d) Representative image of a cell death assay showing autoactive cell death triggered by transient expression of the HREZ D472V MHD motif mutant, but not by wild-type (WT) HREZ.

### HREZ^D472V^ triggers constitutive cell death in *Nicotiana benthamiana*

HREZ carries the canonical CC, NB-ARC and LRR domain architecture, with a conserved MHD motif in the NB-ARC domain (**Fig. 1c**). Specific mutations in the MHD motif can render NLRs autoactive in the absence of pathogen effectors. We generated an HREZ variant by mutating the conserved MHD motif aspartate to valine (D472V) to test whether the variant triggers constitutive cell death. Transient overexpression assays in *N. benthamiana* confirmed that the MHD motif variant, HREZ^D472V^, triggered constitutive cell death, whereas wild-type HREZ remained inactive when expressed alone, consistent with autoinhibition of the receptor in its resting state (**Fig. 1c,d**).

### Activated HREZ assembles into a higher-order complex

Recognition of the ToBRFV MP and the constitutive cell death in HREZ^D472V^ prompted us to examine whether HREZ undergoes oligomerization following activation. To facilitate biochemical analysis of activated HREZ, we generated an HREZ variant carrying substitutions in the N-terminal α1 helix (HREZ^M9E/L13E/I17E^), based on mutations previously shown to abolish cell death without affecting receptor activation or oligomerization in other CC-NLRs (Adachi *et al*., 2019; Madhuprakash *et al*., 2024). C-terminally FLAG-tagged HREZ^M9E/L13E/I17E^ was co-expressed with the C-terminally V5-tagged ToBRFV MP in *N. benthamiana* and analyzed by blue native polyacrylamide gel electrophoresis (BN-PAGE). A prominent high-molecular-weight HREZ complex was detected in the presence of the MP but was absent in the GFP control (**Fig. 2a**). Immunoblotting with an anti-V5 antibody did not detect the MP at the corresponding molecular weight, suggesting that the MP is not stably retained within the activated HREZ complex under the conditions tested.

**Fig. 2.**
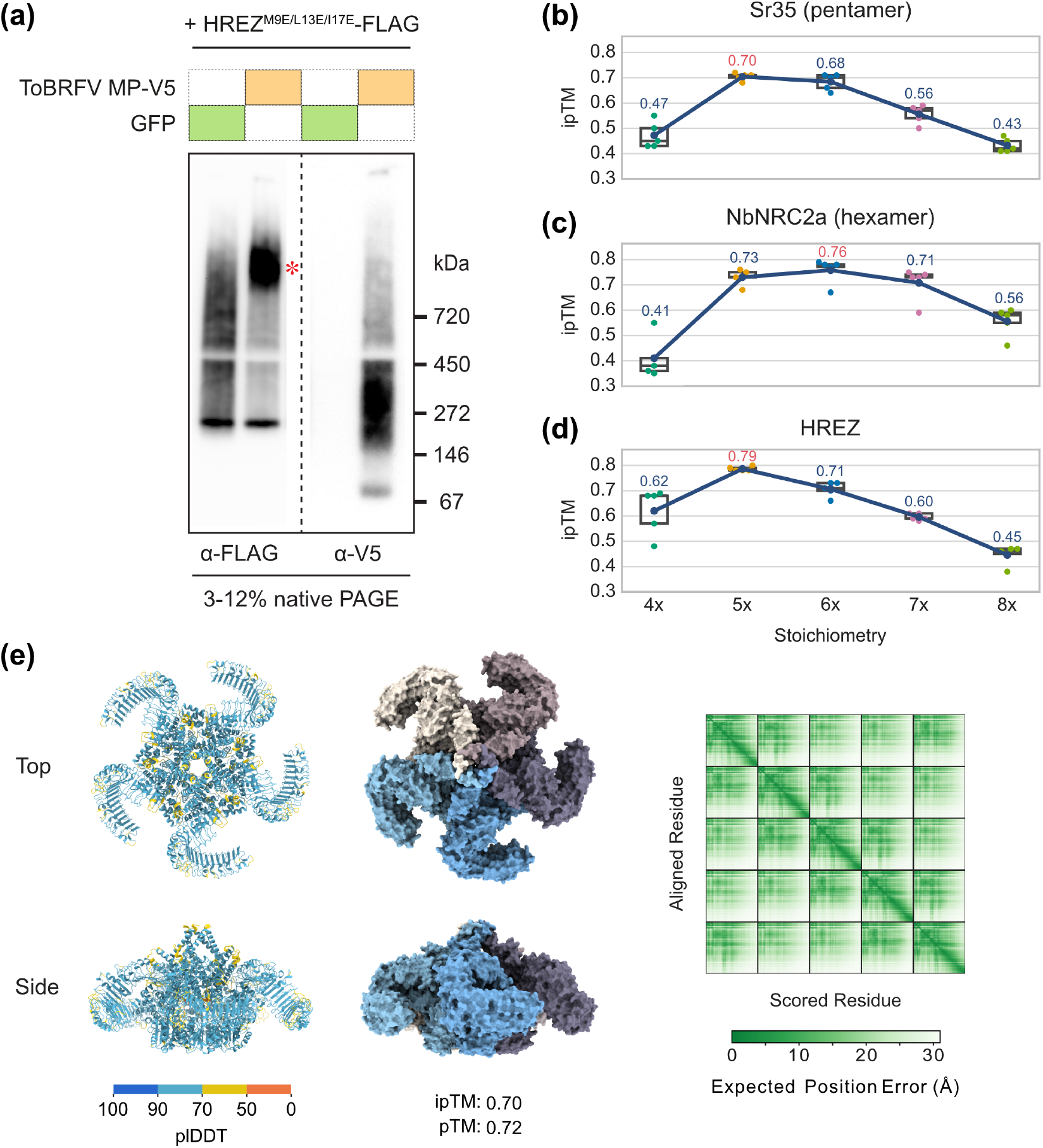
HREZ likely forms a pentameric resistosome upon activation by the ToBRFV MP. (a) Blue native polyacrylamide gel electrophoresis (BN-PAGE) assay showing putative HREZ resistosome formation upon co-expression with the ToBRFV MP. FLAG-tagged HREZ^M9E/L13E/I17E^ variant was co-expressed with either V5-tagged ToBRFV MP or GFP as a control. The red asterisk indicates the putative resistosome band. (b-d) ipTM scores from AlphaFold 3 predictions of Sr35 (b), NbNRC2a (c), and HREZ (d) for different stoichiometries. Amino acid sequences spanning the oligomerizing domains of each NLR protein, from the N terminus to the end of the NB-ARC domain, were used for the predictions. The structures were modelled in the presence of 25 oleic acid molecules using five different seed values, seeds 1 to 5. Sr35 and NbNRC2a are known to form pentameric and hexameric resistosomes, respectively. (e) AlphaFold 3 prediction of the pentameric resistosome structure of full-length HREZ. The structural models are shown coloured by pLDDT confidence scores and by individual protomers. The predicted aligned error (PAE) plot is also shown.

To infer the oligomeric state of activated HREZ, we predicted the oligomeric structures of HREZ alongside those of the experimentally determined pentameric Sr35 and hexameric NbNRC2a resistosomes using AlphaFold 3 (Zhao *et al*., 2022; Förderer *et al*., 2022; Abramson *et al*., 2024; Madhuprakash *et al*., 2024). For each NLR, assemblies containing four to eight protomers were modelled with five different seeds using sequences spanning the N terminus to the end of the NB-ARC domain, with 25 oleic acid molecules included in each prediction as a proxy for cellular membranes. For both Sr35 and NbNRC2a, the experimentally determined oligomeric state yielded the highest interface predicted TM (ipTM) scores among the stoichiometries tested, supporting the ability of this modelling approach to recover known oligomeric states (**Fig. 2b,c**; **Table S1**). Among the HREZ models, the pentamer consistently yielded the highest ipTM scores across independent predictions (**Fig. 2d**). We additionally modelled a full-length HREZ pentamer in the absence of oleic acid molecules, which adopted a funnel-shaped structure (**Fig. 2e**). Collectively, these results demonstrate activation-dependent HREZ oligomerization and support a pentameric model for the activated HREZ complex.

### HREZ and Tm-2^2^ belong to phylogenetically distinct CC-NLR lineages

HREZ has been described to confer resistance to ToBRFV, but its evolutionary relationship to the previously characterized tobamovirus-resistance NLR Tm-2^2^ has remained unclear. To determine the phylogenetic position of HREZ, we reconstructed a maximum-likelihood phylogeny using 4,657 NB-ARC domain sequences from 12 Solanaceae and three outgroup species (*Coffea arabica, Ipomoea nil*, and *Arabidopsis thaliana*) (**Fig. 3**; **Table S2**). HREZ clustered within a CC-NLR lineage distinct from that containing Tm-2^2^. In addition, neither HREZ nor Tm-2^2^ belonged to the highly expanded NRC network (Wu *et al*., 2017) (**Fig. 3**). Together with their shared responsiveness to tobamovirus MPs, these phylogenetic relationships support independent evolution of tobamovirus MP recognition in the HREZ and Tm-2^2^ CC-NLR lineages.

**Fig. 3.**
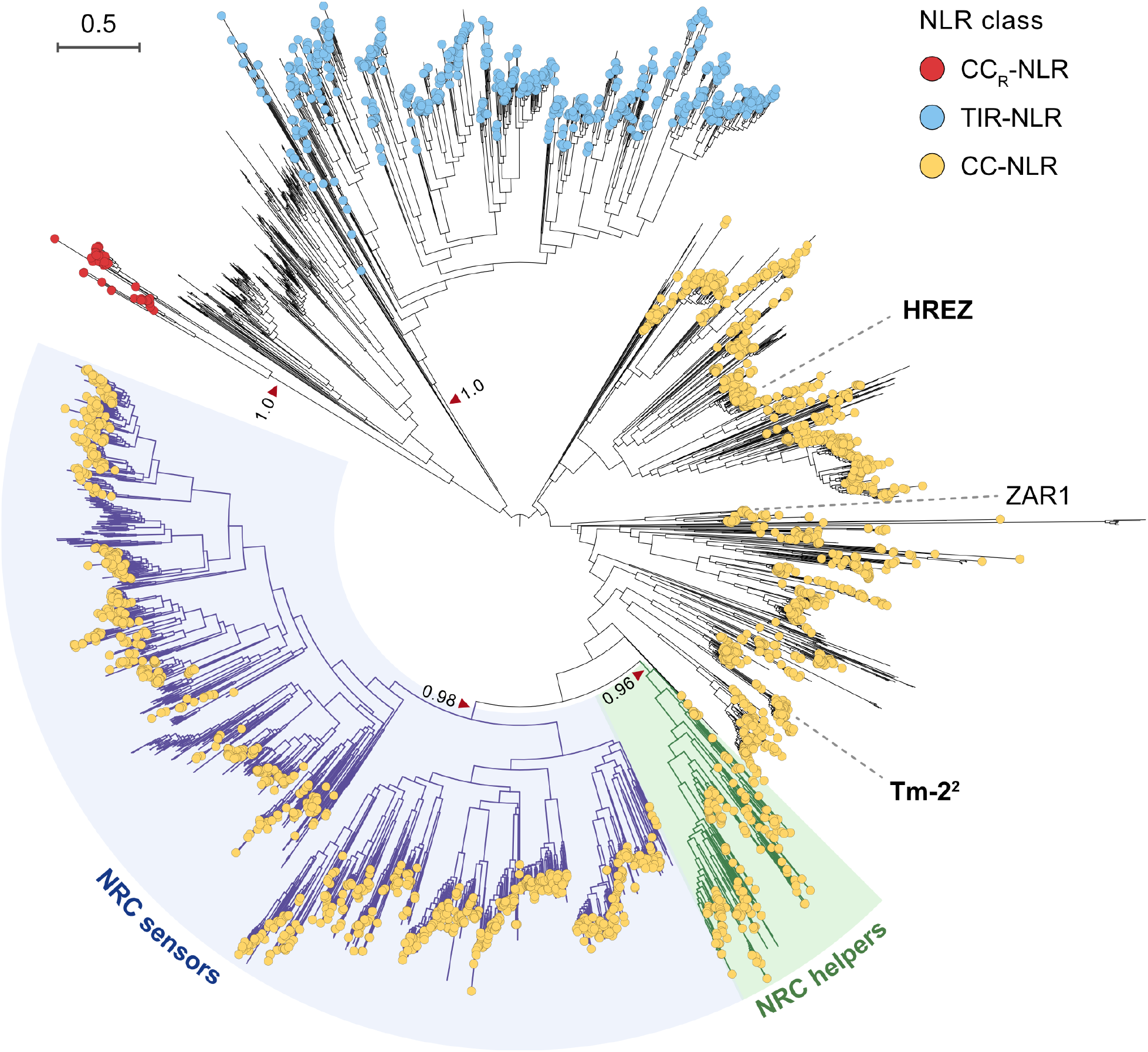
HREZ and Tm-2^2^ belong to phylogenetically distinct CC-NLR lineages. Phylogenetic tree of 4,657 deduplicated NLR NB-ARC amino acid sequences from HREZ, Tm-2^2^, RefPlantNLR reference proteins (Kourelis *et al*., 2021), and NLRtracker-predicted NLRs from selected plant genomes, including 12 Solanaceae species and three outgroup species, *Coffea arabica, Ipomoea nil*, and *Arabidopsis thaliana* (Toghani & Kamoun, 2024). The maximum-likelihood tree was inferred using VeryFastTree (Piñeiro & Pichel, 2024). Tip circles indicate NLR class assignments based on domain-architecture annotations: CC_R_-NLR, red; TIR-NLR, blue; and CC-NLR, yellow. Colours were assigned only to sequences with a domain architecture supporting a unique CC_R_-NLR, TIR-NLR, or CC-NLR classification, defined by the presence of RPW8, TIR, or CC together with NB-ARC and LRR domains. Shaded clades indicate NRC helper and sensor lineages.

### HREZ and Tm-2^2^ exhibit distinct recognition spectra for tobamovirus movement proteins

Given that HREZ is phylogenetically distinct from Tm-2^2^, we next asked whether this evolutionary divergence is reflected in their recognition spectra for tobamovirus MPs. To address this, we reconstructed a phylogeny of 32 diverse MPs and then selected eight MPs representing the phylogenetic diversity within the family *Virgaviridae* (**Fig. 4a**; **Table S3**). Using transient co-expression assays in *N. benthamiana*, we assessed cell death and ion leakage (**Fig. 4b,c**). HREZ responded to the MPs of TMV, ToBRFV, ToMV and Pepper mild mottle virus (PMMoV). In contrast, Tm-2^2^ responded to the MPs of TMV, ToMV, PMMoV, Paprika mild mottle virus (PaMMV), Turnip vein-clearing virus (TVCV) and Cucumber green mottle mosaic virus (CGMMV). Tm-2^2^ also showed a weak cell-death phenotype in response to the ToBRFV MP, although this response did not result in a significant increase in ion leakage (**Fig. 4b,c**). Neither receptor responded to the MP of Tobacco rattle virus (TRV), a member of the genus *Tobravirus* whose MP was the most phylogenetically divergent among those tested. These results demonstrate partially overlapping but distinct recognition spectra for HREZ and Tm-2^2^.

**Fig. 4.**
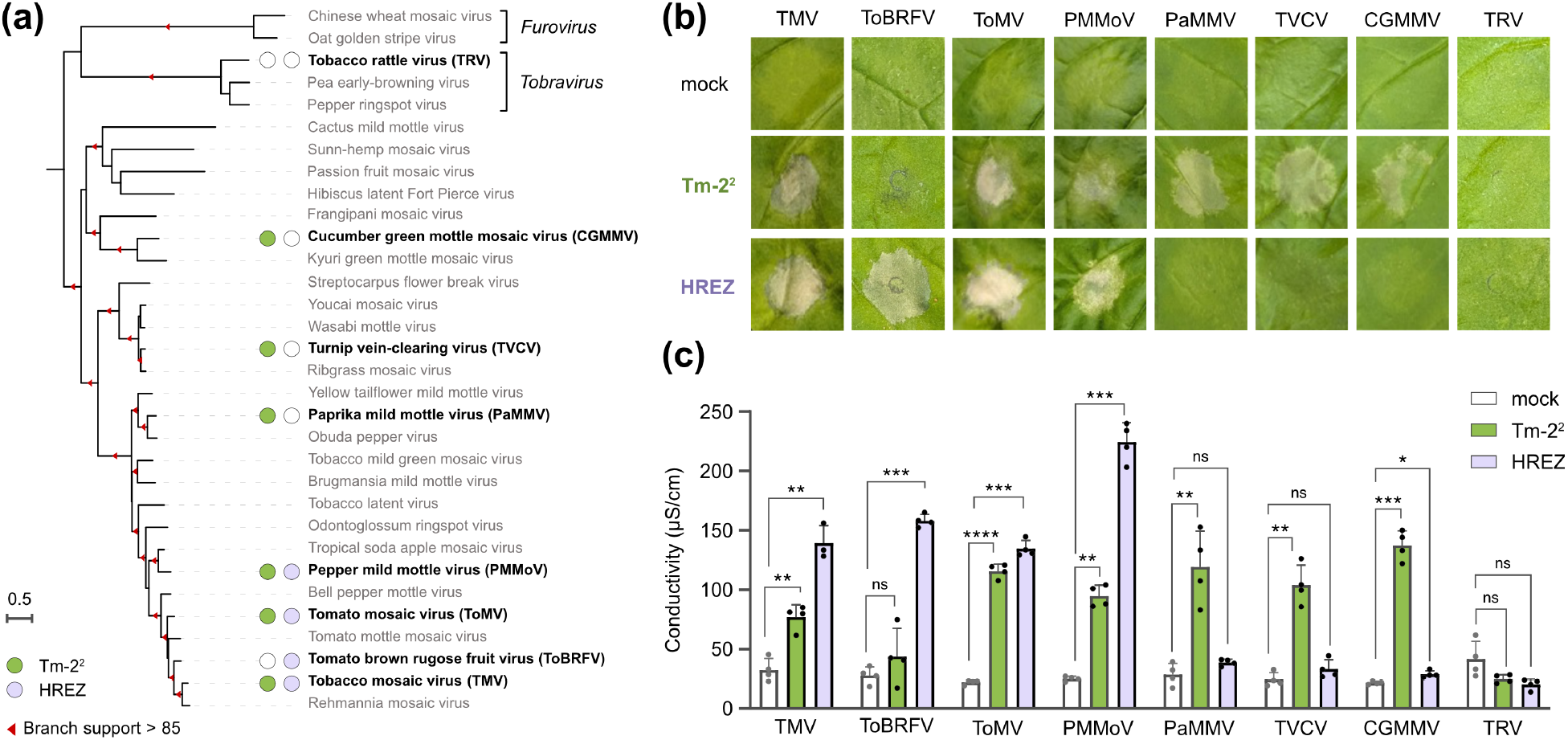
Comparative analysis of HREZ and Tm-2^2^ immune responses to MPs from the family *Virgaviridae* reveals distinct recognition spectra. (a) Phylogenetic tree of 32 *Virgaviridae* MPs, including the eight MPs tested in this study. The tree was inferred from a codon-based nucleotide sequence alignment using IQ-TREE 2 (Minh *et al*., 2020). MPs recognized by either Tm-2^2^ or HREZ are indicated in the tree. (b) Representative images of *N. benthamiana* leaf sections co-agroinfiltrated with each of eight MPs together with Tm-2^2^, HREZ, or *Agrobacterium* carrying an empty vector as a mock control. Leaves were photographed 5 days after infiltration. (c) Ion leakage assay after co-agroinfiltration of the eight MPs with mock, Tm-2^2^, or HREZ. Leaf discs were collected 22 h post-infiltration, and conductivity was measured after 6 h of incubation in double-distilled water. Data are from four biological replicates, with four leaf discs per replicate. Statistical analysis was performed using Brown-Forsythe and Welch ANOVA. Asterisks indicate significant differences compared with the mock control: ****P < 0.0001, ***P < 0.001, **P < 0.01, and *P < 0.05.

### HREZ-recognized tobamovirus movement proteins promote HREZ oligomerization

We next investigated whether different tobamovirus MPs recognized by HREZ promote higher-order complex assembly. BN-PAGE analysis revealed that the MPs from TMV, ToBRFV, ToMV and PMMoV each induced the formation of a prominent high-molecular-weight HREZ complex, whereas the MPs from PaMMV, TVCV, CGMMV and TRV did not induce detectable HREZ oligomerization (**Fig. 5**). The MPs that promoted HREZ oligomerization were the same as those that elicited HREZ-dependent cell death and increased ion leakage. Furthermore, immunoblotting with an anti-V5 antibody failed to detect TMV, ToBRFV, ToMV or PMMoV MPs within the high-molecular-weight HREZ complex, despite their ability to trigger oligomerization. These results suggest that HREZ-mediated immune activation by MPs is associated with HREZ oligomerization, and that the assembled complex does not stably retain these MPs under the conditions tested.

**Fig. 5.**
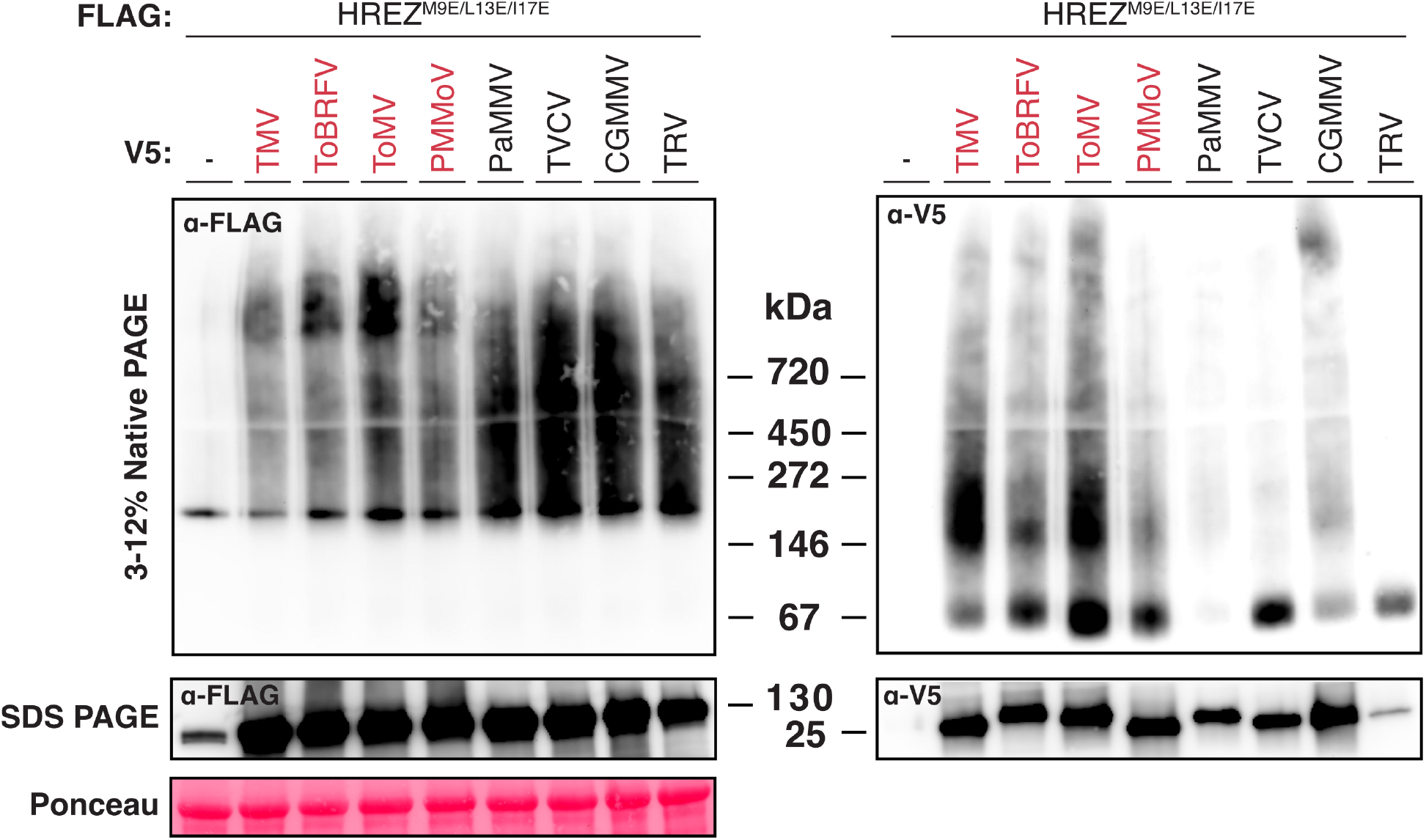
HREZ-recognized tobamovirus movement proteins promote HREZ oligomerization. Blue native polyacrylamide gel electrophoresis (BN-PAGE) assay showing higher-order HREZ resistosome formation upon co-expression with tobamovirus MPs. FLAG-tagged HREZ^M9E/L13E/I17E^ mutant was co-expressed with each of the eight V5-tagged MPs tested in Fig. 4.

## Discussion

The rapid global emergence of ToBRFV has highlighted the need for new genetic sources of resistance that complement existing resistance genes such as *Tm-2*^*2*^. Here, we show that the *S. habrochaites* CC-NLR HREZ recognizes the ToBRFV MP, triggering HREZ-dependent cell death and higher-order receptor assembly, with AlphaFold 3 modelling supporting a pentameric resistosome model (**Fig. 1, 2**). Phylogenetic analyses revealed that HREZ and Tm-2^2^ belong to distinct CC-NLR lineages but display partially overlapping recognition spectra for tobamovirus MPs (**Fig. 3, 4**). Notably, the MPs recognized by HREZ were also those that promoted HREZ oligomerization, linking recognition specificity to activation-dependent higher-order assembly (**Fig. 5**). Together, these findings reveal convergent recognition of tobamovirus MPs by phylogenetically distinct CC-NLR lineages, while showing that HREZ and Tm-2^2^ provide distinct and complementary recognition spectra.

Our results support an executor function for HREZ, but whether HREZ functions as a singleton NLR remains unresolved. Its autoactivity, activation-dependent oligomerization, predicted pentameric resistosome, and phylogenetic placement outside the NRC network are all consistent with a singleton mode of action (**Fig. 1–3**). However, the activating MPs were not detectably retained within the assembled HREZ complex by BN-PAGE (**Fig. 2a, 5**). This differs from structurally characterized singleton NLR resistosomes such as Sr35 and ZAR1, in which the activating effector or receptor-associated intermediate remains incorporated in the assembled complex (Wang *et al*., 2019; Zhao *et al*., 2022; Förderer *et al*., 2022). One possibility is that HREZ itself recognizes tobamovirus MPs through a weak or transient interaction that is not maintained after resistosome assembly. Such a lack of stable incorporation of an upstream activator into the assembled resistosome has precedent in the Rx/NRC helper (Ahn *et al*., 2023b; Contreras *et al*., 2023b; Toghani *et al*., 2026a). Alternatively, because HREZ was isolated from the wild tomato species *S. habrochaites* and our assays were performed in *N. benthamiana*, an endogenous *N. benthamiana* immune component could perceive the MP and activate HREZ downstream. Determining whether HREZ itself mediates MP recognition and resolving the structure of the activated complex will be important for defining its mode of action.

Intriguingly, *HREZ*-mediated resistance is known to be temperature sensitive, with reduced effectiveness at elevated temperatures (Jewehan *et al*., 2022c; Verweij *et al*., 2026). Recent work suggests that elevated temperature can impair NLR oligomerization, raising the possibility that HREZ thermosensitivity may similarly arise at the level of resistosome assembly (Grech-Baran *et al*., 2025). Our assays for MP-induced HREZ cell death and oligomerization in *N. benthamiana* provide a tractable system for future work on HREZ thermosensitivity (**Fig. 1, 2a**). Defining this mechanism may facilitate the development of HREZ variants with improved temperature tolerance.

We compared the recognition spectrum of HREZ to Tm-2^2^ (**Fig. 4**). Tm-2^2^ has been known to recognize MPs from multiple tobamoviruses, although the breadth of this recognition has remained poorly defined (Weber & Pfitzner, 1998; Chen *et al*., 2017; Hak *et al*., 2023; Tettey *et al*., 2023). Within the panel examined here, Tm-2^2^ responded to a phylogenetically diverse range of MPs and exhibited a broader recognition spectrum than HREZ (**Fig. 4**). Tm-2^2^ also showed a weak cell-death phenotype in response to the ToBRFV MP, although this response was not accompanied by a significant increase in ion leakage (**Fig. 4b,c**). This observation may be compatible with recent evidence that Tm-2^2^ retains attenuated recognition of the ToBRFV MP despite the ability of ToBRFV to overcome *Tm-2*^*2*^-mediated resistance (Zhang *et al*., 2026). Although Tm-2^2^ has been shown to associate with tobamovirus MPs and to undergo activation-dependent self-association (Chen *et al*., 2017; Wang *et al*., 2020), how it recognizes such a diverse range of MPs and translates their recognition into receptor activation remains unclear.

Plant immune receptors can independently evolve to recognize the same pathogen effector or activity, as exemplified by RPS5/PBR1 and Ptr1/ZAR1 (Carter *et al*., 2019; Ahn *et al*., 2023a). HREZ and Tm-2^2^ represent a distinct example of such convergence in antiviral immunity: although they belong to separate CC-NLR lineages, both recognize tobamovirus MPs but exhibit distinct, partially overlapping recognition spectra (**Fig. 4**). This suggests that convergence on the same viral target does not necessarily entail convergence in recognition specificity. Tm-2^2^ recognized a broader range of MPs than HREZ, whereas HREZ strongly recognized the ToBRFV MP, which evades effective *Tm-2*^*2*^-mediated resistance. Thus, HREZ should not be viewed simply as a replacement for Tm-2^2^, but rather as a complementary source of tobamovirus resistance, providing a rationale for combining phylogenetically distinct resistance sources to broaden protection against current and emerging tobamoviruses.

## Materials and Methods

### Plasmid constructions

The plasmids and expression constructs used in this study are listed in **Table S4**, together with their corresponding vector backbones, tags, and relevant sequence information.

### Plant materials

*N. benthamiana* plants were grown in soil in a light- and temperature-controlled chamber at 25°C under a 16-h photoperiod. The leaves of four-to five-week-old plants were used for agroinfiltration.

### Cell death assays

For the ToBRFV protein screening and HREZ autoactivity assays, HREZ, Pikobody^α-GFP^, and the indicated ToBRFV proteins were transiently expressed in *N. benthamiana* leaves by *Agrobacterium tumefaciens*-mediated infiltration. Cultures of *A. tumefaciens* strain GV3101 harbouring the indicated binary vectors were grown overnight at 28°C, resuspended in infiltration buffer containing 10 mM MgCl_2_, 10 mM MES, and 150 μM acetosyringone (pH 5.6), and adjusted to OD_600_ = 0.3. For co-expression assays, cultures carrying the indicated NLR and viral protein constructs were mixed at a 1:1 ratio before infiltration. For the autoactivity assay, HREZ or HREZ^D472V^ was infiltrated individually at OD_600_ = 0.3. For the comparative MP recognition assays of HREZ and Tm-2^2^, cultures of *A. tumefaciens* strain EHA105 harbouring the indicated binary vectors were grown overnight at 28°C, resuspended in the same infiltration buffer, and adjusted to OD_600_ = 0.5. *Agrobacterium* suspensions carrying HREZ, Tm-2^2^, or an empty-vector control were co-infiltrated with suspensions carrying the indicated MP constructs. For both assays, *Agrobacterium* suspensions were infiltrated into the abaxial side of *N. benthamiana* leaves using a needleless syringe. Cell death phenotypes were assessed and photographed 5 days after infiltration.

### Protein extraction

For biochemical analysis of HREZ oligomerization, *N. benthamiana* leaves were agroinfiltrated with *A. tumefaciens* strain GV3101 carrying FLAG-tagged HREZ^M9E/L13E/I17E^ together with either the indicated V5-tagged MP constructs or a GFP control. Bacterial cultures were grown overnight at 28°C, resuspended in the same infiltration buffer used for the cell death assays, and adjusted to OD_600_ = 0.3. Three days after infiltration, six leaf discs were harvested per sample and homogenized using a Geno/Grinder tissue homogenizer. Ground tissue was resuspended in GTMN buffer [10% (v/v) glycerol, 50 mM Tris-HCl (pH 7.5), 5 mM MgCl_2_, and 50 mM NaCl] supplemented with 10 mM dithiothreitol (DTT), 1× protease inhibitor cocktail (Sigma-Aldrich), and 0.2% (v/v) NP-40 substitute (Sigma-Aldrich). Samples were incubated on ice for 10 min with brief vortexing every 2 min to facilitate extraction. Lysates were centrifuged at 5,000 × g for 20 min at 4°C, and the resulting supernatants were collected for BN-PAGE and SDS-PAGE analyses.

### BN-PAGE and SDS-PAGE analysis

For BN-PAGE, clarified protein extracts were diluted according to the manufacturer’s instructions with NativePAGE™ 5% G-250 Sample Additive, 4× NativePAGE™ Sample Buffer, and deionized water. Samples were loaded onto 3–12% Bis-Tris NativePAGE™ gels (Invitrogen) and electrophoresed alongside SERVA Native Marker (SERVA). Following electrophoresis, proteins were transferred to polyvinylidene difluoride (PVDF) membranes using NuPAGE™ Transfer Buffer and the Trans-Blot Turbo Transfer System (Bio-Rad), according to the manufacturer’s instructions. Membranes were fixed in 8% acetic acid for 15 min, rinsed with water, and air-dried. To visualize native protein standards, membranes were reactivated with ethanol prior to immunoblotting.

For SDS-PAGE, protein samples were mixed with SDS loading buffer and denatured at 72 °C for 10 min. Following centrifugation at 5,000 × g for 3 min, the supernatants were loaded onto 4–20% Mini-PROTEAN^®^ TGX™ precast gels (Bio-Rad) and electrophoresed alongside the PageRuler™ Plus Prestained Protein Ladder (Thermo Scientific). Proteins were transferred to PVDF membranes using the Trans-Blot Turbo Transfer System with the supplied transfer buffer, following the manufacturer’s instructions.

### Immunoblotting and detection

Membranes were blocked in 5% (w/v) non-fat dry milk in Tris-buffered saline containing 0.01% Tween-20 (TBS-T) for 1 h at room temperature. Blots were then incubated overnight at 4 °C with horseradish peroxidase (HRP)-conjugated primary antibodies diluted 1:5,000 in 5% milk/TBS-T. The following antibodies were used: anti-GFP (B-2, Santa Cruz Biotechnology), anti-V5 (V5-10, Sigma-Aldrich), and anti-FLAG (M2, Sigma-Aldrich).

Signals were detected using Pierce™ ECL Western Blotting Substrate (32106, Thermo Fisher Scientific), with up to 50% SuperSignal™ West Femto Maximum Sensitivity Substrate (34095, Thermo Fisher Scientific) added when enhanced sensitivity was required. Chemiluminescence was imaged using either an ImageQuant LAS 4000 or an ImageQuant 800 system (GE Healthcare Life Sciences). Membrane loading controls were visualized by staining with Ponceau S (Sigma-Aldrich) or Ponceau 4R (AG Barr).

### Electrolyte leakage assay

Constructs harbouring each MP under the 35S promoter (p35S) were mixed at a 1:1 ratio with either *p35S:Tm-2*^*2*^, *p35S:HREZ*, or an empty-vector control, and the mixtures were agroinfiltrated into *N. benthamiana* leaves (n ≥ 4) at OD_600_ = 0.25. Four leaf discs (1 cm diameter) were cut from each leaf and washed for 10–30 min in distilled water, 22 h after agroinfiltration. The leaf discs were then carefully transferred into a tube containing 4 mL of distilled water and placed on a low-speed orbital shaker. Electrical conductivity was measured after 6 h using a benchtop conductivity meter.

### AlphaFold 3 predictions

Based on NLRtracker outputs, amino acid sequences spanning the N terminus to the end of the NB-ARC domain were extracted using a previously published script (Toghani *et al*., 2025). Using the AlphaFold 3 web server (https://alphafoldserver.com/), assemblies containing four to eight protomers were modelled for each NLR with 25 oleic acid molecules included in each prediction. Five independent predictions were performed using different seed values (1–5). In addition, a full-length HREZ pentamer was modelled without oleic acid. Predicted structures were visualized using UCSF ChimeraX (Meng *et al*., 2023). The AlphaFold 3 prediction dataset generated in this study has been archived in Zenodo (https://doi.org/10.5281/zenodo.22046208).

### Phylogenetic analysis of HREZ and Tm-2^2^

The dataset comprised HREZ, the experimentally validated RefPlantNLR collection including Tm-2^2^ (Kourelis *et al*., 2021), and NLRtracker annotations from 15 RefSeq plant proteomes representing 12 Solanaceae and three outgroup species (Toghani & Kamoun, 2024) (**Table S2**). Only NLRtracker records designated Status = NLR were retained, and multiple NB-ARC domains from the same protein were treated as separate records. Sequences 250–400 amino acids in length were retained from the merged set of 8,540 NB-ARC sequences using SeqKit v2.10.0 (Shen *et al*., 2024). Exact amino acid sequence duplicates were removed, yielding 4,657 unique sequences. The NB-ARC sequences were aligned using FAMSA v2.4.1 with the option -refine_mode on (Gudyś *et al*., 2026). The alignment was trimmed using ClipKIT v2.4.1 in gappy mode with the options ‘-m gappy -g 0.9’ (Steenwyk *et al*., 2020). A maximum-likelihood tree was inferred with VeryFastTree v4.0.5 using default protein settings (Piñeiro & Pichel, 2024). The tree was midpoint-rooted using ETE v4.3.0 (Huerta-Cepas *et al*., 2016) and visualized and annotated with iTOL v6 (Letunic & Bork, 2024). NLR classes were assigned from the simplified full-length domain-architecture labels generated by NLRtracker. A sequence was assigned to the CC_R_-NLR, TIR-NLR, or CC-NLR class when its label contained exactly one of the corresponding strings: RNL, TNL, or CNL. The NRC helper and sensor clades were delineated using curated tip lists anchored by the corresponding RefPlantNLR reference proteins. The commands, processed input datasets, and output files are available in the GitHub repository (https://github.com/YuSugihara/HREZ_NLR_MP_phylogeny).

### Phylogenetic analysis of *Virgaviridae* movement proteins

Thirty-two representative viruses classified within the family *Virgaviridae* according to the International Committee on Taxonomy of Viruses (ICTV) were selected, comprising 27 members of the genus *Tobamovirus*, three of *Tobravirus*, and two of *Furovirus* (Adams *et al*., 2017) (**Table S3**). The corresponding full-length amino acid sequences and coding sequences (CDSs) were extracted using their NCBI protein accessions. Full-length MP amino acid sequences were aligned using MAFFT with the options ‘--globalpair --maxiterate 1000’ (Katoh & Standley, 2013). The corresponding CDSs were then threaded onto the protein alignment using the ‘thread_dna’ function of PhyKIT (Steenwyk *et al*., 2021), producing a codon-aware alignment. A maximum-likelihood tree was inferred from this nucleotide alignment using IQ-TREE v2.3.6 with ModelFinder (Kalyaanamoorthy *et al*., 2017; Minh *et al*., 2020). The GTR+F+R4 substitution model was selected according to the Bayesian information criterion, and branch support was assessed using 1,000 replicates of the SH-like approximate likelihood-ratio test (‘-alrt 1000’). The commands, processed input datasets, and output files are available in the GitHub repository (https://github.com/YuSugihara/HREZ_NLR_MP_phylogeny).

## Data availability

The AlphaFold 3 prediction dataset generated in this study has been archived in Zenodo (https://doi.org/10.5281/zenodo.22046208). The commands, processed input datasets, sequence alignments, and phylogenetic tree files used for the phylogenetic analyses of HREZ/Tm-2^2^ and *Virgaviridae* movement proteins are available in the GitHub repository (https://github.com/YuSugihara/HREZ_NLR_MP_phylogeny).

## Author contributions

Conceptualization: Y.S., J.M., Methodology: Y.S., H.H., N.L., J.M.A., Z.S., J.M., Investigation: Y.S., H.H., N.L., J.M.A., Z.S., J.M., Formal analysis: Y.S., H.H., N.L., J.M.A., Z.S., J.M., Resources: Y.S., Z.S., J.M., Writing – original draft: Y.S., J.M. Writing – review & editing: Y.S., Z.S., J.M., Visualization: Y.S., Z.S., J.M., Supervision: Y.S., Z.S., J.M., Project administration: Y.S., Z.S., J.M., Funding acquisition: Y.S., Z.S., J.M.

## Acknowledgements

We thank Prof. Sophien Kamoun (The Sainsbury Laboratory) for his critical input and support. We also thank Dr. Jiorgos Kourelis (Imperial College London) for his valuable suggestions on cloning the HREZ and ToBRFV MP genes. We acknowledge the use of the colour palettes created by Chenxin Li for figure preparation.

